# Domain-general brain regions do not track linguistic input as closely as language-selective regions

**DOI:** 10.1101/076240

**Authors:** Idan Blank, Evelina Fedorenko

## Abstract

Language comprehension engages a cortical network of left frontal and temporal regions. Activity in this network is language-selective, showing virtually no modulation by non-linguistic tasks. In addition, language comprehension engages a second network consisting of bilateral frontal, parietal, cingulate, and insular regions. Activity in this “Multiple Demand (MD)” network scales with comprehension difficulty, but also with cognitive effort across a wide range of non-linguistic tasks in a domain-general fashion. Given the functional dissociation between the language and MD networks, their respective contributions to comprehension are likely distinct, yet such differences remain elusive. Critically, given that each network is sensitive to some linguistic features, prior research has assumed – implicitly or explicitly – that both networks track linguistic input closely, and in a manner consistent across individuals. Here, we used fMRI to directly test this assumption by comparing the BOLD signal time-courses in each network across different people listening to the same story. Language network activity showed fewer individual differences, indicative of closer input tracking, whereas MD network activity was more idiosyncratic and, moreover, showed lower reliability within an individual across repetitions of a story. These findings constrain cognitive models of language comprehension by suggesting a novel distinction between the processes implemented in the language and MD networks.

**Significance Statement:** Language comprehension recruits both language-specific mechanisms and domain-general mechanisms that are engaged in many cognitive processes. In the human cortex, language-selective mechanisms are implemented in the left-lateralized “core language network”, whereas domain-general mechanisms are implemented in the bilateral “Multiple Demand (MD)” network. Here, we report the first direct comparison of the respective contributions of these networks to naturalistic story comprehension. Using a novel combination of neuroimaging approaches we find that MD regions track stories less closely than language regions. This finding constrains the possible contributions of the MD network to comprehension, contrasts with accounts positing that this network has continuous access to linguistic input, and suggests a new typology of comprehension processes based on their extent of input tracking.

## Introduction

A key desideratum for a theory of language comprehension is to specify the division of linguistic labor across distinct cognitive mechanisms. Insofar as distinct mechanisms are implemented in separable neural populations, one promising approach to advance such theories is to functionally characterize brain regions or networks that are engaged in comprehension and establish their respective roles. Indeed, high-level language processing recruits several large-scale networks, each exhibiting a unique functional profile. Among these, the network most critical to comprehension is the “core language network”, a set of left frontal and temporal regions. This network is robustly engaged in language processing (e.g., Binder et al., 1997; Jung-Beeman, 2005; Fedorenko et al., 2010; Menenti et al., 2011) across languages (e.g., Sebastian et al., 2011), presentation modalities (Chee et al., 1999; Buchweitz et al., 2009; Fedorenko et al., 2010; Braze et al., 2011; Vagharchakian et al., 2012), and developmental experiences (Neville et al., 1998; Bedny et al., 2011). It exhibits sensitivity to linguistic features such as lexico-semantic information and syntactic structure (Keller et al., 2001; Fedorenko et al., 2012b; Bautista and Wilson, 2016; Blank et al., 2016) but, critically, shows virtually no engagement in non-linguistic tasks and is therefore language-selective (Fedorenko et al., 2011; for a review, see: Fedorenko and Varley, 2016).

In addition, language comprehension engages the “multiple demand (MD)” network (Duncan, 2010), consisting of frontal, parietal, cingulate, and insular regions bilaterally. Activity in this network is sensitive to comprehension difficulty, increasing in response to e.g., temporary ambiguity and non-local syntactic dependencies (for a review, see: Fedorenko, 2014). However, this network similarly scales its response with cognitive effort across a wide range of non-linguistic tasks (Duncan and Owen, 2000; Miller and Cohen, 2001; Braver et al., 2003; Dosenbach et al., 2006; Cole and Schneider, 2007; Fedorenko et al., 2013; Hugdahl et al., 2015) and is therefore domain-general.

The strikingly different functional profiles of the language and MD networks are evident not only in task-based neuroimaging studies but also in relatively unconstrained, task-free paradigms where these networks show unsynchronized activity fluctuations during naturalistic cognition (Blank et al., 2014). A similar double dissociation is reported by neuropsychological studies: damage to the language network leads to language impairments (Broca, 1861/2006; Dax, 1863; Wernicke, 1874/1969; Geschwind, 1970; Bates et al., 2003) but leaves other high-level cognitive functions largely intact (Fedorenko and Varley, 2016); whereas damage to the MD network impairs executive functions (Luria, 1966/2012; Fuster, 1989; Woolgar et al., 2010), but can leave comprehension mostly unimpaired (for a review, see: Fedorenko, 2014). Finally, language and executive functions follow different developmental trajectories (Fedorenko, 2014). These converging findings, of course, do not imply that domain-general executive functions do not play a role in language processing (for discussions, see Fedorenko, 2014; Geranmayeh et al., 2014). Indeed, a number of studies have provided causal evidence linking executive control to language (Wiener et al., 2004; Fridriksson et al., 2006; Amici et al., 2007; Murray, 2012). However, they provide evidence that the language and MD networks support distinct computations and their contributions to comprehension fundamentally differ.

Nonetheless, the precise nature of the respective contributions remains elusive, as most prior neuroimaging studies of comprehension have not couched their findings in terms of the functional distinction between the language and MD networks. To the extent that available accounts do draw this distinction, however implicitly, they suggest that the two networks differ in either the input features that they each track or the operations they each engage in while processing such input (e.g., Novick et al., 2005; Thompson-Schill et al., 2005; Hickok and Poeppel, 2007; Friederici, 2012; Hagoort, 2013; Fedorenko, 2014). Critically, the various postulated roles of these two networks appear to depend on continuous access to the unfolding input: such access underlies any suggestion that neural activity co-varies with a certain linguistic feature, e.g., lexical information, compositional structure, local ambiguity or parsing difficulty. However, this assumption – which is crucial for understanding the contributions of the two networks to language processing – has not been empirically evaluated.

Here, we use fMRI to directly test this assumption. Namely, we measure activity fluctuations in the language and MD networks during story comprehension and estimate how tightly coupled those fluctuations are to the story. Consistent with current views, one might hypothesize that both networks exhibit equally close tracking of linguistic input. Alternatively, one network might track the input less closely than the other, in which case the space of operations such a network could support would be importantly constrained. Such a finding would indicate that the contributions of the two networks to comprehension differ more fundamentally than is presently assumed.

## Materials and Methods

Below, we outline and motivate our methodology. Specifically, we describe a novel combination of existing approaches that is designed to meet four criteria: (i) high functional resolution for identifying brain networks; (ii) a naturalistic paradigm suitable for studying comprehension in all its richness (cf. traditional task-based paradigms); (iii) direct comparisons of brain networks for valid statistical inferences; and (iv) reproducibility of results.

In order to evaluate the extent of input tracking in the language and MD networks, we first must define the cortical regions of interest that constitute these networks. In this process, we must account for the fact that individual brains are highly variable in the mapping of high-level cognitive functions onto macro-anatomical landmarks. This variability, evident in the temporal cortex (Jones and Powell, 1970; Gloor, 1997; Wise et al., 2001) and especially in the frontal cortex (Amunts et al., 1999; Tomaiuolo et al., 1999; Chein et al., 2002) where language and MD regions lie side-by-side (Fedorenko et al., 2012a), renders anatomical localization precarious (Juch et al., 2005; Poldrack, 2006; Fischl et al., 2008; Tahmasebi et al., 2011; Frost and Goebel, 2012). For these reasons, we similarly cannot rely on functional localization at the level of an entire sample using group-based analyses (Saxe et al., 2006; Fedorenko and Kanwisher, 2009). Therefore, we functionally localize language and MD regions individually in each participant. This approach allows us to pool data from the same functional regions across participants even when those regions do not align well spatially.

Following functional localization, we evaluate how closely the language and MD networks track linguistic input during naturalistic comprehension. Our interest in naturalistic input is threefold: first, some brain regions respond more reliably to richly structured natural input compared to experimentally controlled input (Hasson et al., 2010). Second, unlike traditional experimental paradigms which often require participants to perform artificial tasks on linguistic materials, naturalistic comprehension more closely approximates language processing “in the wild”, where the primary goal is the extraction of meaning. Therefore, this “task free” paradigm provides an important complementary approach for evaluating the contributions of the MD regions to comprehension, especially given that these regions operate in a task-dependent manner (Miller and Cohen, 2001; Sreenivasan et al., 2014; D’Esposito and Postle, 2015). And third, naturalistic comprehension requires all aspects of the input to be combined into a single rich representation, unlike experimental stimuli and tasks that focus on particular linguistic features and have lower ecological validity. Therefore, we record the BOLD signal fluctuations of language and MD regions while participants passively listen to stories, where the only explicit task is to comprehend the story’s content.

Following Lerner et al. (2011), we reasoned that if a given network closely tracked the story such that fluctuations in its BOLD signal were stimulus-locked, then its signal time-course would be similar across participants and would thus show a high Inter-Subject Correlation (ISC) (Hasson et al., 2004). Hence, we use ISC as an index of input tracking. Critically, ISC is a “model-free” measure: instead of testing how well signal time-courses can be explained by certain pre-specified, hypothesis-driven predictors, each participant’s empirical data serve as the model compared against the data from the other participants.

This data-driven method has been successfully used to demonstrate that broad cortical swathes do track stories to significant extents (Wilson et al., 2008; Lerner et al., 2011; Honey et al., 2012; Regev et al., 2013; Silbert et al., 2014; Schmälzle et al., 2015), proposing a neural correlate of “shared understanding” across individuals (Hasson et al., 2012). Nevertheless, prior studies have measured ISCs in a voxel-wise fashion, whereby brains were first anatomically aligned and, then, each stereotaxic location served in turn as a basis for comparing signal time-courses across participants. Relating the resulting cortical topology of ISCs to the topology of known functional brain networks could then proceed only through “reverse inference” (Poldrack, 2006). Moreover, voxel-wise comparisons across participants rely on the invalid assumption that a given anatomical location has a common function across individuals. To relax this assumption, here we augment the ISC framework by comparing signal time-courses across regions that are functionally defined. This allows us to focus on, and compare between, language and MD regions, such that we can tie our findings to the wealth of prior literature characterizing the response profiles of those networks.

In addition, we augment the statistical approach adopted in early studies of ISCs by directly testing the correlations in the language network against those in the MD network. Such an explicit comparison between networks allows for more nuanced inferences compared to those licensed when each network is separately tested against a null baseline and differences across networks are indirectly inferred (cf. Lerner et al., 2011, for the latter) (Nieuwenhuis et al., 2011).

Finally, we demonstrate that our results are reproducible, by reporting two replications of our main, story comprehension experiment: the first is a direct replication with a subset of the original stories; the second is a conceptual replication with a new, even more naturalistic story.

### Participants

Fifty participants between the ages of 18 and 47, recruited from the MIT student body and the surrounding community, were paid for participation. Two participants were removed from the analysis due to poor quality of the functional localizer data and three more were removed due to poor segmentation of their anatomical scan. Of the remaining 45 participants (30 females; mean age 23.5, SD 4.8), 19 were tested in the main experiment, 13 in the first replication and 19 in the second replication (the first and third groups were partially overlapping). In addition, 15 of these participants were tested in a control experiment (described below): these included 8 participants from the main experiment, 2 from the first replication, and one who participated in both the main experiment and the second replication. Forty-one participants were right-handed (based on the Edinburgh Handedness Inventory; Oldfield, 1971), and the remaining four left-handed participants had a left-lateralized language network (for motivation to include left-handers in cognitive neuroscience research, see Willems et al., 2014). All participants were native English speakers and gave informed consent in accordance with the requirements of MIT’s Committee on the Use of Humans as Experimental Subjects (COUHES).

### Design, stimuli and procedure

#### Language localizer task

The task used to localize the language network is described in detail in Fedorenko et al. (2010). Briefly, we used a reading task contrasting sentences and lists of unconnected, pronounceable nonwords (Figure 1a) in a standard, deterministic blocked design with a counterbalanced order across runs (for timing parameters, see Table 1). Stimuli were presented one word / nonword at a time. For the first ten participants only, each trial ended with a memory probe and they had to indicate, via a button press, whether or not that probe had appeared in the preceding sequence of words / nonwords. The remaining participants instead read the materials passively (we included a button-pressing task at the end of each trial, to help participants remain alert). Importantly, this localizer has been shown to generalize across task manipulations: the sentences > nonwords contrast robustly activates the fronto-temporal language network regardless of the task (Fedorenko et al., 2010). The regions identified by this contrast engage in a broad range of linguistic processes including (but not limited to) lexico-semantic processes and combinatorial syntactic and semantic processes (Fedorenko et al., 2012b; Blank et al., 2016; Fedorenko et al., 2016; Fedorenko et al., 2017). Moreover, this localizer identifies the same regions that are localized with a broader contrast, between recorded natural speech and its acoustically-degraded version (Scott et al., 2016).

**Figure 1.**
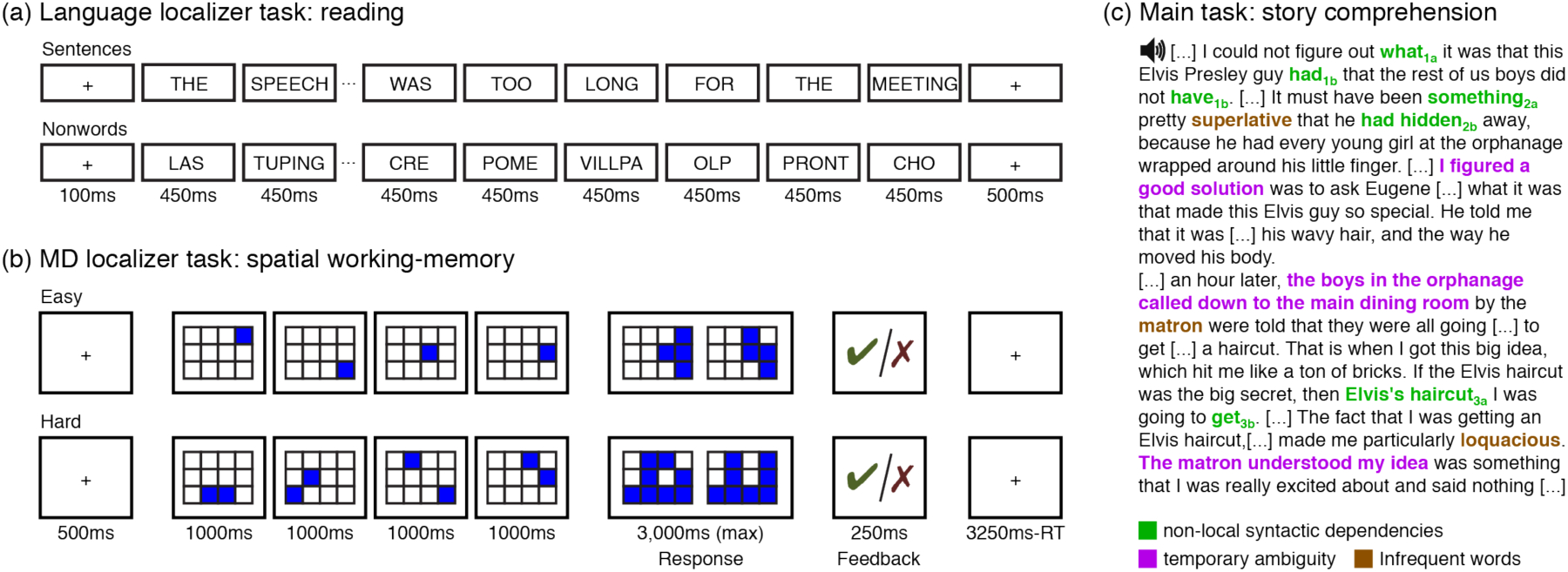
Experimental tasks. (a) The reading task used to localize language regions, based on the critical contrast sentences > nonwords. (b) The spatial working-memory task used to localize MD regions, based on the critical contrast hard > easy. (c) An excerpt from a story used in the main comprehension experiment. Linguistic phenomena that increase processing difficulty and have been shown to recruit the MD network, but are naturally infrequent, were edited into the text. These include non-local syntactic dependencies (green; words in this relation have subscripts with the same number but different letters); temporary ambiguity (purple), where a likely initial parse is later revealed to be wrong; and low-frequency words (brown).

**Table 1.** Timing parameters for the different versions of the language localizer task.

|  | Version |  |  |
| --- | --- | --- | --- |
|  | A | B | C |
| Number of participants | 35 | 5 | 5 |
| Task: Passive Reading or Memory? | PR | M | M |
| Words / nonwords per trial | 12 | 12 | 12 |
| Trial duration (ms) | 6,000 | 6,000 | 6,000 |
| Fixation | 100 | --- | --- |
| Presentation of each word / nonword | 450 | 350 | 350 |
| Fixation | 500 | 300 | 300 |
| Memory probe | --- | 1,000 | 1,000 |
| Fixation | --- | 500 | 500 |
| Trials per block | 3 | 3 | 3 |
| Block duration (s) | 18 | 18 | 18 |
| Blocks per condition (per run) | 8 | 8 | 6 |
| Conditions | Sentences<br>Nonwords | Sentences<br>Nonwords | Sentences<br>Nonwords<br>Word-lists<sup>1</sup> |
| Fixation block duration (s) | 14 | 18 | 18 |
| Number of fixation blocks | 5 | 5 | 4 |
| Total run time (s) | 358 | 378 | 396 |
| Number of runs | 2 | 2 | 2-3 |

<sup>1</sup> Used for the purposes of another experiment; see (Fedorenko et al., 2010)

#### MD localizer task

Regions of the MD network were localized using a spatial working-memory task contrasting a hard version with an easy version (Figure 1b). On each trial (8s), participants saw a 3×4 grid and kept track of eight (hard version) or four (easy version) randomly generated locations that were sequentially flashed two at a time or one at a time, respectively (1s per flash). Then, participants indicated their memory for these locations in a two-alternative, forced-choice paradigm via a button press (3s total). Feedback was immediately provided upon choice (or lack thereof) (250ms). Hard and easy conditions were presented in a standard blocked design (4 trials in a 32s block, 6 blocks per condition per run) with a counterbalanced order across runs. Each run included 4 blocks of fixation (16s each) and lasted a total of 448s. Thirty-nine participants completed 1-2 runs of the localizer. The remaining six participants either provided poor-quality data (5 participants) or were not run on this task (1 participant). For this latter group, MD regions were localized with data from the language localizer task, using the (reverse) nonwords > sentences contrast. Both the hard > easy contrast and the nonwords > sentences contrast have been previously demonstrated to robustly and reliably identify the MD network (Fedorenko et al., 2013) (these participants did not differ from the rest of the sample in the dependent variables; see Table 2a).

**Table 2.**
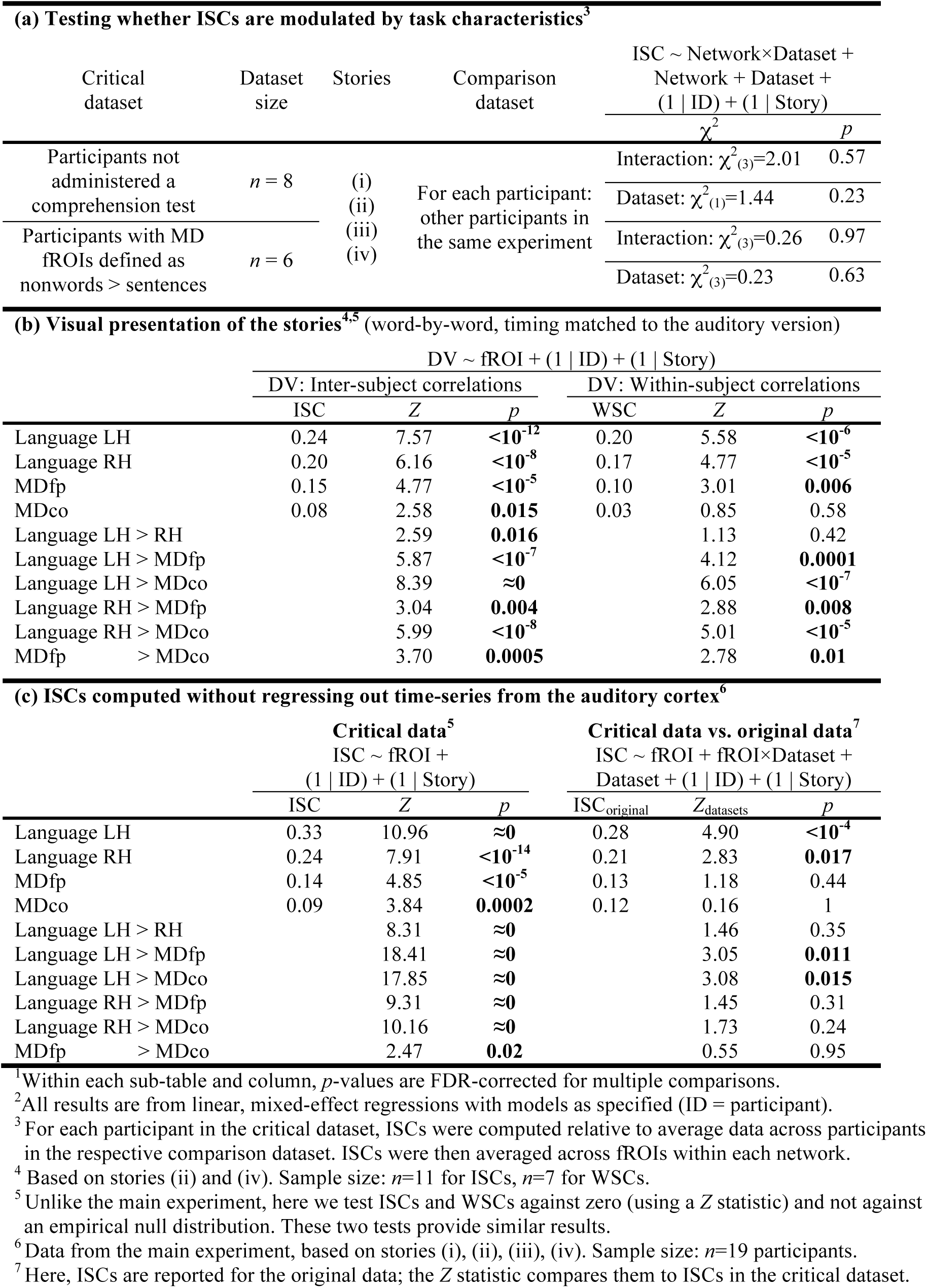
The patterns of story tracking across the language and MD networks generalize across changes in the experimental setup and analysis stream^1,2^.

#### Story comprehension task

In the main experiment, each subject listened to 1-4 stories (one story: *n*=7; two: *n*=3; three: *n*=2; four: *n=7*; duration: 270s-364s) over scanner-safe headphones (Sensimetrics, Malden, MA). Stories were constructed based on publicly available fairy tales and short stories:

i. “The Legend of the Bradford Boar” (by E. H. Hopkinson; unedited version: www.make4fun.com/stories/British-short-story/3917-The-Legend-of-the-Bradford-Boar-by-E-H-Hopkinson)
ii. “Aqua; or the Water Baby” (by Kate Douglas Wiggin; unedited version:fullreads.com/literature/aqua-or-the-water-baby/)
iii. “The King of the Birds” (by The Brothers Grimm; unedited version:www.apples4theteacher.com/holidays/bird-day/short-stories/the-king-of-the-birds.html)
iv. “Elvis Died at the Florida Barber College” (by Roger Dean Kiser; unedited version: www.eastoftheweb.com/short-stories/UBooks/ElvDie.shtml).

These stories were edited to include a variety of linguistic phenomena that have been shown to increase processing difficulty in numerous prior behavioral sentence processing studies and which recruit the MD network (Figure 1c; see also Supplementary Materials). As a result of these edits, comprehension difficulty was robustly modulated across each story. Namely, self-paced reading times in a separate sample (*n=*181 participants) were reliably predicted by measures of linguistic complexity (Shain et al., 2016). Moreover, in these stories, some measures of complexity influenced online behavior more robustly than in studies that have used unedited texts, plausibly because the relevant linguistic phenomena do not naturally occur with sufficiently high frequency (Collins, 1996; Roland et al., 2007; Futrell et al., 2015). Further, even though the stories in the current experiments were presented via the auditory rather than visual modality, we still expect them to successfully modulate processing difficulty because reading-time effects generalize to on-line listening (Ferreira et al., 1996; Waters and Caplan, 2001) (see also Table 2b for evidence that our neuroimaging results generalize to visual story presentation).

In the first replication, participants listened to stories (i) and (iii) used in the main experiment (these data were originally collected for the purpose of a separate experiment; participants also listened to the other two stories, but performed a simultaneous, unrelated task during those trials). In the second replication, participants listened to an autobiographical story (“Pie-man,” told by Jim O’Grady) recorded at a live storytelling event (“The Moth” storytelling event, NYC). This story (duration: 420s) did not undergo linguistic editing and was thus even more naturalistic than the previous stories. Each story started and ended with 16s seconds of fixation (and music, for the Pie-man story) that were not analyzed.

To test the reliability of signal time-courses in the language and MD networks, participants in the control experiment listened to the same stories twice, either within the same scanning session (approximately one hour apart, *n*=7) or in separate sessions (6.5-21.5 months apart, *n*=8) (4 participants listened to the same story twice within the same session and then, once more, in a separate session).

After each story, participants answered 6-12 comprehension questions that required attentive listening (i.e., could not have been answered correctly based on common knowledge). For the main experiment and the first replication, participants answered two-alternative forced-choice (2AFC) questions via a button press while in the scanner. For the second replication, participants filled in a 4AFC questionnaire after the scanning session. For eight participants, answers to these questions were not recorded due to equipment malfunction (these participants did not differ from the rest of the sample in the dependent variables; see table 2a). The remaining 37 participants demonstrated good comprehension, with a negatively skewed accuracy distribution (mode=100%, median=87.5%, semi-interquartile range=12.85%).

### Data acquisition and preprocessing

#### Data acquisition

Whole-brain structural and functional data were collected on a whole-body 3 Tesla Siemens Trio scanner with a 32-channel head. T1-weighted structural images were collected in 176 axial slices with 1mm isotropic voxels (repetition time (TR) = 2,530ms; echo time (TE) = 3.48ms). Functional, blood oxygenation level-dependent (BOLD) data were acquired using an EPI sequence with a 90° flip angle and using GRAPPA with an acceleration factor of 2; the following parameters were used: thirty-one 4mm thick near-axial slices acquired in an interleaved order (with 10% distance factor), with an in-plane resolution of 2.1mm × 2.1mm, FoV in the phase encoding (A ≫ P) direction 200mm and matrix size 96mm × 96mm, TR = 2000ms and TE = 30ms. The first 10s of each run were excluded to allow for steady state magnetization.

#### Spatial preprocessing

Data preprocessing was carried out with SPM5 (using default parameters, unless specified otherwise) and supporting, custom MATLAB scripts. Preprocessing of anatomical data included normalization into a common space (Montreal Neurological Institute (MNI) template), resampling into 2mm isotropic voxels, and segmentation into probabilistic maps of the gray matter, white matter (WM) and cerebrospinal fluid (CSF). Preprocessing of functional data included motion correction (realignment to the mean image using 2^nd^-degree b-spline interpolation), normalization (estimated for the mean image using trilinear interpolation), resampling into 2mm isotropic voxels, smoothing with a 4mm FWHM Gaussian filter and high-pass filtering at 200s.

#### Temporal preprocessing

Additional preprocessing of data from the story comprehension runs was carried out using the CONN toolbox (Whitfield-Gabrieli and Nieto-Castanon, 2012) with default parameters, unless specified otherwise. Five temporal principal components of the BOLD signal time-courses extracted from the WM were regressed out of each voxel’s time-course; signal originating in the CSF was similarly regressed out. Six principal components of the six motion parameters estimated during offline motion correction were also regressed out, as well as their first time derivative. Next, the residual signal was bandpass filtered (0.008–0.09 Hz) to preserve only low-frequency signal fluctuations (Cordes et al., 2001). This filtering did not influence the results reported below.

#### Participant-specific functional localization of language and MD networks Modeling localizer data

For each localizer task, a standard mass univariate analysis was performed in SPM5 whereby a general linear model estimated the effect size of each condition in each experimental run. These effects were each modeled with a boxcar function (representing entire blocks) convolved with the canonical Hemodynamic Response Function (HRF). The model also included first-order temporal derivatives of these effects, as well as nuisance regressors representing entire experimental runs and offline-estimated motion parameters. The obtained beta weights were then used to compute the functional contrast of interest: for the language localizer, sentences > nonwords, and for the MD localizer, hard > easy (or nonwords > sentence for 6 participants; see Design, stimuli and procedure).

#### Defining functional regions of interest (fROIs)

Language and MD fROIs were defined based on functional contrast maps from the localizer experiments. These maps were first restricted to include only gray matter voxels by excluding voxels that were more likely to belong to either the white matter or the cerebrospinal fluid based on SPM’s probabilistic segmentation of the participant’s structural data.

Then, fROIs in the language network were defined using group-constrained, participant-specific localization (Fedorenko et al., 2010). For each participant, the map of the sentences > nonwords contrast was intersected with binary masks that constrained the participant-specific language network to fall within areas where activations for this contrast are relatively likely across the population. These masks are based on a group-level representation of the contrast obtained from a previous sample. We used 8 such masks in the left-hemisphere, including regions in the posterior, mid-posterior, mid-anterior and anterior temporal lobe, as well as in the middle frontal gyrus, the inferior frontal gyrus and its orbital part (Figure 2a). These masks were mirror-projected onto the right-hemisphere to create 8 homologous masks (the masks cover significant parts of the cortex, so their mirrored version is likely to encompass the right-hemisphere homologue of the left-hemisphere language network, despite possible hemispheric asymmetries in their precise locations). In each of the resulting 16 masks, a participant-specific language fROI was defined as the top 10% of voxels with the highest contrast values. This top *n*% approach ensures that fROIs can be defined in every participant and that their sizes are the same across participants, allowing for generalizable results (Nieto-Castañón and Fedorenko, 2012).

**Figure 2.**
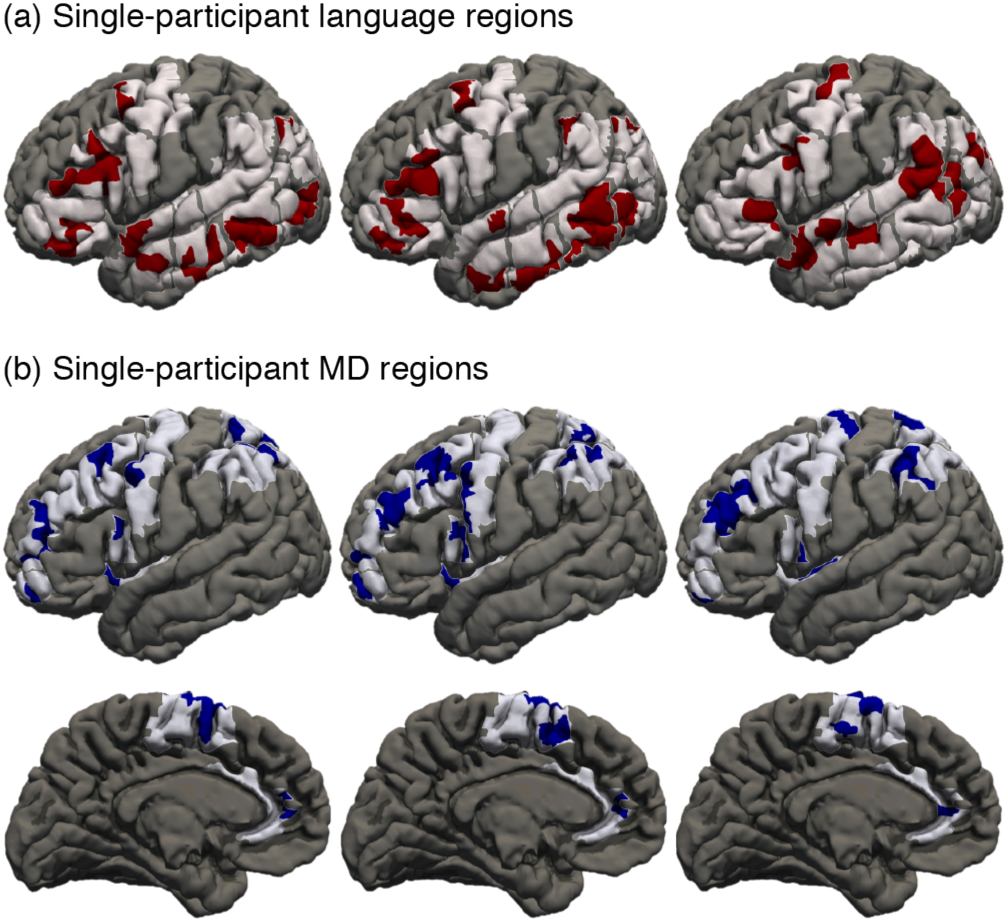
Functional regions of the language and MD networks. (a) LH language regions in 3 individual participants are shown in dark red. These regions were localized with a reading task (see Figure 1a). These regions were constrained to fall within eight broad areas where activations for this task are common across the population, shown in light pink. These areas were defined based on group-level data from a previous sample (Fedorenko et al., 2010). (b) LH MD regions of the same 3 participants are shown in dark blue. These regions were localized with a spatial working-memory task (see Figure 1b). These regions were constrained to fall within nine broad areas where activations for this localizer are common across the population, shown in light blue. These areas were anatomically defined (Fedorenko et al., 2013). Apparent overlap between language and MD fROIs is illusory and due to projection onto the cortical surface.

fROIs in the MD network were similarly defined (using the “top 10%” approach) based on the hard > easy contrast in the spatial working-memory game. Here, instead of using binary masks based on group-level functional data, we used anatomical masks (Tzourio-Mazoyer et al., 2002; see Fedorenko et al., 2013; Blank et al., 2014). Nine masks were used in each hemisphere, including regions in the middle frontal gyrus and its orbital part, the opercular part of the inferior frontal gyrus, the precental gyrus, the posterior and inferior parts of the partietal lobe, the insula, and supplementary motor area and the cingulate cortex (Figure 2b). Based on prior findings (Dosenbach et al., 2006; Dosenbach et al., 2007; Nomura et al., 2010; Power et al., 2011; Mantini et al., 2013), we grouped the resulting fROIs into two functionally distinct sub-networks: fronto-parietal (first 5 masks) and cingulo-opercular (last 3 masks). Similar results were obtained when fROIs were instead grouped by hemisphere. (We note that functional masks derived for the MD network based on 197 participants significantly overlapped with the anatomical masks; we chose to use the anatomical masks in order to maintain comparability between our functional data and data from previous studies that have used these masks).

Any voxels that were identified by both the language and the MD localizer were excluded from analysis (this procedure did not influence the results). Language fROIs had a median overlap of 0 voxels with the MD network (inter-quartile range: 2.1%). MD fROIs also had a median overlap of 0 voxels with the language network (inter-quartile range: 1.6%). The resulting fROIs had an average size of 247±77 voxels in the language network, and 212±111 voxels in the MD network.

### Critical analysis: inter-subject correlations (ISCs)

#### Computing ISCs

For each participant and fROI, BOLD signal time-courses recorded during story comprehension were extracted from each voxel beginning 6 seconds following the onset of the story (to exclude an initial rise in the hemodynamic response relative to fixation, which could increase ISCs). These time-courses were first temporally *z*-scored in each voxel and then averaged across voxels. Next, to ensure that the resulting signal time-course reflected the tracking of high-level linguistic information and not low-level sensory information, we removed from it any variance that was explained by activity in the auditory cortex. Specifically, the signal was regressed against signals extracted from anatomically defined regions around the postero-medial and antero-lateral sections of Heschl’s gyrus bilaterally (Tzourio-Mazoyer et al., 2002) (this regression did not affect the pattern of results reported here; see Table 2c). Finally, for each story, participant, and fROI we computed an ISC value, namely, Pearson’s moment correlation coefficient between the residual time-course and the corresponding average residual time-course across the remaining participants (Lerner et al., 2011). ISCs were Fisher-transformed prior to statistical testing in order to improve normality (Silver and Dunlap, 1987).

#### Statistical testing

In each fROI, ISCs were then tested for significance against an empirical null distribution based on 1,000 simulated signal time-courses that were generated by phase-randomization of the original data (Theiler et al., 1992). Namely, we generated null distributions for individual participants, fit each distribution with a Gaussian, and analytically combined the resulting parameters across participants. The true ISCs, also averaged across participants, were then *z*-scored relative to these empirical parameters and converted to one-tailed *p*-values.

Critically, ISCs were compared across networks using a linear, mixed-effects regression (Barr et al., 2013) implemented with the “lme4” package in R. In each experiment, ISCs across all fROIs, participants and stories were modeled with a fixed effect of fROI and random intercepts for participant and story. The fixed effect estimates were combined across fROIs within each functional network (LH language, RH language, fronto-parietal MD, and cingulo-opercular MD) and were pairwise compared to each other using the “multcomp” package in R. Hypotheses were two-tailed for the first experiment and one-tailed for the replications and control analyses. In each experiment, *p*-values are reported following False Discovery Rate (FDR) correction for multiple comparisons (Benjamini and Yekutieli, 2001).

For all findings based on linear, mixed-effects regression analyses, similar results were obtained when data for each participant were first averaged across fROIs within each network and pairwise network comparisons (across participants) were then tested using exact permutation tests (Gill, 2007). Therefore, our results are independent of assumptions regarding data normality.

### Control analysis: within-subject correlations (WSCs)

#### Computing WSCs

For each participant who listened to the same story on two separate trials, we computed a within-subject correlation (WSC) value for each fROI by correlating the signal time-courses across the two trials. The resulting correlations were Fisher-transformed.

Note that unlike ISCs, which compare the signal from one participant to an average signal across all other participants, WSCs compare two single-trial signals. Consequently, the two measures are not directly comparable: despite the fact that WSCs are not contaminated by inter-individual variability and should thus be higher than ISCs, ISCs will *de facto* be higher because signal averaging removes a lot of noise from the data. In order to make ISCs comparable to WSCs we therefore computed “pairwise ISCs”: for each participant and fROI, we correlated the signal time-course separately with each of the corresponding, individual signal time-courses of the other participants, Fisher-transformed the resulting correlation values, and averaged them.

#### Statistical tests

Prior to these analyses, we tested whether WSCs in the within-session and across-session datasets differed from each other. To this end, we performed a linear, mixed-effects regression analysis that modeled individual WSCs for all fROIs, participants and stories with a fixed effect of the interaction between fROI and dataset, random intercepts for participant and story, and a random slope for dataset varying by participant (this model was chosen because a fuller model failed to converge). Pairwise contrasts tested whether WSCs in each network were stronger across sessions than within a session. These two groups did not differ from each other in their network WSCs. Therefore, these two sets of data were modeled together in the critical analyses: here, WSCs were compared across networks using the same model that was used to test ISCs, modeling individual WSCs for all fROIs, participants and stories

A similar approach was used for comparing WSCs to pairwise-ISCs. Here, contrasts tested whether pairwise differences between networks observed with WSCs were distinct from those observed with ISCs.

## Results

### Correlations of network activity across individuals listening to the same story

ISC data are presented in Figure 3. Across stories in the main experiment, the left-hemisphere (LH) language network showed the highest ISCs (Fisher transformed *r*=0.280), stronger than ISCs in the right-hemisphere (RH) language network (*r*=0.210; Cohen’s *d*=0.73, *z*=6.25, *p*<10^−9^), the fronto-parietal MD (MDfp) network (*r*=0.136; *d*=1.07, *z*=14.12, *p≈*0) and the cingulo-opercular MD (MDco) network (*r*=0.117; *d*=1.32, *z*=13.51, *p≈*0). The RH language network, in turn, showed higher ISCs than both the MDfp network (*d*=1.07, *z*=7.27, *p*<10^−11^) and the MDco network (*d*=1.04, *z*=7.72, *p*<10^−13^). The two MD networks did not differ from each other (*d*=1.80, *z*=1.70, *p*=0.218). The difference between the LH language network and the two MD networks was also observed for each story separately.

**Figure 3.**
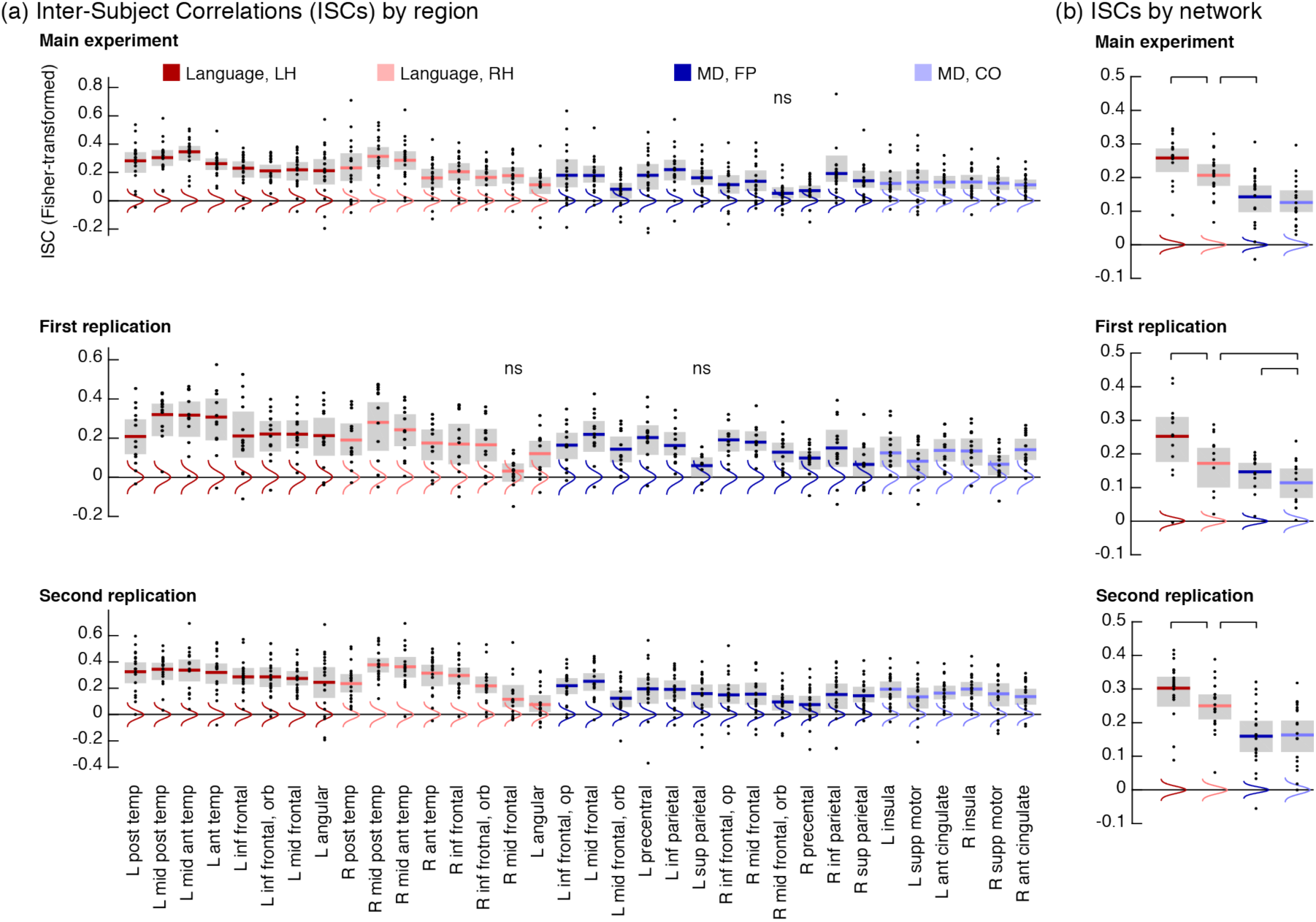
ISCs during story comprehension in the language and MD networks. (a) ISC (Fisher-transformed) for each brain region. Black dots are individual data points. Thick, colored horizontal lines show the average ISCs across participants. Gray rectangles show 95% confidence intervals of these average ISCs (empirically derived using 1,000 permutations). Colored vertical curves show Gaussian fits to empirical null distributions against which average ISCs can be tested (ns, non-significant results at a threshold of 0.05; FDR-corrected). Regions are grouped into 4 functional networks, indicated by color. Across experiments, a replicable pattern emerges where ISCs are stronger in language regions (red) than in MD regions (blue). (b) Mean ISCs within each functional network, same conventions as in (a). Black, horizontal lines connect pairs of networks that significantly differ from one another (in each pair, the left ISC is greater than the right ISCs and all ISCs that are further to the right). L – left; R – right; Post – posterior; Temp – temporal; Mid – middle; Ant – anterior; Inf – inferior; Orb – orbital; Op – opercular; Sup – superior; Supp – supplementary.

In both replication experiments, we again found that ISCs in the LH language network (replication 1: *r*=0.252; replication 2: *r*=0.303) were stronger than in the RH language network (*r*=0.172, *d*=0.90, *z*=5.62, *p*<10^−7^; *r*=0.250, *d*=0.77, *z*=3.35, *p*=0.001), the MDfp network (*r*=0.147, *d*=1.06, *z*=8.09, *p*<10^−15^; *r*=0.160, *d*=1.29, *z*=9.95, *p≈*0) and the MDco network (*r*=0.114, *d*=1.33, *z*=8.95, *p≈*0; *r*=0.163, *d*=1.34, *z*=8.20, *p*<10^−15^). ISCs in the RH language network were somewhat stronger than ISCs in the MDfp network (*d*=0.46, *z*=1.93, *p*=0.066; *d*=0.82, *z*=6.28, *p*<10^−9^) and stronger than ISCs in the MDco network (*d*=0.70, *z*=3.74, *p*<0.001; *d*=0.83, *z*=5.10, *p*<10^−7^). The two latter networks reliably differed from each other only in the first replication (*d*=0.53, *z*=2.28, *p*<0.033).

Across these three experiments, we find that signals in the language and MD networks differ in their ISCs and, thus, in the percentage of variance they share across individuals. To further interpret these findings we computed an “upper bound” on ISCs, reflecting the highest values that could be expected in our measurements; namely, we computed ISCs in low-level auditory regions (see Materials and Methods) that track sensory input very closely (Lerner et al., 2011). Combining data across experiments, these auditory ISCs are estimated at *r*=0.450. Thus, signals in the LH language network (*r*=0.287) share 40.8% of this “maximum shareable variance” across individuals; signals in the RH language network (*r*=0.216) share 23%, whereas signals in the MDfp network (*r*=0.153) and MDco network (*r*=0.134) share only 11.6% and 8.8%, respectively. Importantly, however, almost all ISCs – even those in MD regions – are significantly greater than expected by chance (Figure 3). Therefore, even domain-general MD regions track stories to a non-trivial extent in spite of doing so substantially and reliably more weakly than the language regions.

Is it possible that other sub-regions of the MD network, not identified by our localizer, track the stories more strongly? To test this possibility, we computed traditional, voxel-based ISCs (based on anatomical alignment of individual brains) and identified, within each mask of the MD network, the voxels that showed the highest ISCs during one story. These voxels served as “alternative fROIs”, and we estimated the strength of their ISCs using independent data from another story. The resulting ISCs were weaker than those reported above (Table 3), and the same finding held in “alternative fROIs” identified in the language network. Critically, compared to the original fROIs, the alternative fROIs responded less robustly to the language and MD localizers (responses in the original fROIs were obtained from runs of the localizers that were held-out during fROI definition). For instance, alternative fROIs in the MDco network did not respond differentially to the hard and easy versions of the spatial working memory task; and alternative fROIs in the RH language network did not respond differentially to sentences and nonwords (Table 3). These decreased functional signatures are likely caused by inter-individual variability in the precise anatomical locations of the language and MD regions, such that a given voxel might belong to a certain network in some participants but not others. Therefore, with no means for establishing functional (rather than anatomical) correspondence across individual brains in areas that lie outside of our localizer-defined fROIs, we do not find evidence for close linguistic tracking of input anywhere in the MD network.

**Table 3.** Functional profiles of “alternative” fROIs defined as the top 10% of voxels in each mask showing the highest ISCs (computed based on anatomical alignment across individual brains)^1^.

| DV | Within dataset (new / original): $DV \sim fROI + (1 \vert ID)$ | | | | | | | |
| --- | --- | --- | --- | --- | --- | --- | --- | --- |
| | Across datasets: $DV \sim fROI + Dataset + fROI \times Dataset + (1 + Dataset \vert ID)$ | | | | | | | |
|  | Language LH |  | Language RH |  | MDfp |  | MDco |  |
|  | new | original | new | original | new | original | new | original |
| ISC<sup>2</sup> | 0.22 | 0.28 | 0.18 | 0.21 | 0.11 | 0.15 | 0.09 | 0.11 |
|  | Z=16.2 | Z=13.4 | Z=13.5 | Z=10.1 | Z=8.2 | Z=7.6 | Z=6.3 | Z=5.0 |
| | $p \approx 0$ | $p \approx 0$ | $p \approx 0$ | $p \approx 0$ | $p < 10^{-15}$ | $p < 10^{-13}$ | $p < 10^{-9}$ | $< 10^{-5}$ |
| | $Z=4.78, p < 10^{-5}$ | | $Z=2.38, p=0.03$ | | $Z=4.05, p < 10^{-4}$ | | $Z=1.5, p=0.21$ | |
| Language localizer<sup>3</sup>: sentences > nonwords | 0.17 | 0.63 | -0.01 | 0.21 | -0.24 | -0.38 | -0.03 | -0.13 |
|  | Z=4.4 | Z=12.0 | Z=-0.4 | Z=4.0 | Z=-7.0 | Z=-7.7 | Z=-0.7 | Z=-2.4 |
| | $p < 10^{-4}$ | $p \approx 0$ | $p=1$ | $p=10^{-4}$ | $p < 10^{-11}$ | $p < 10^{-12}$ | $p=0.77$ | $p=0.03$ |
| | $Z=11.9, p \approx 0$ | | $Z=5.9, p < 10^{-7}$ | | $Z=-3.9, p < 10^{-3}$ | | $Z=-2.3, p=0.04$ | |
| MD localizer<sup>3</sup>: hard > easy | -0.03 | -0.12 | 0.06 | 0.01 | 0.37 | 0.95 | 0.04 | 0.52 |
|  | Z=-0.4 | Z=-1.4 | Z=0.8 | Z=0.09 | Z=5.22 | Z=10.5 | Z=0.5 | Z=5.3 |
| | $p=1$ | $p=0.48$ | $p=0.97$ | $p=1$ | $p < 10^{-6}$ | $p \approx 0$ | $p=1$ | $p < 10^{-6}$ |
| | $Z=1.4, p=0.42$ | | $Z=0.7, p=0.94$ | | $Z=8.4, p \approx 0$ | | $Z=5.9, p < 10^{-7}$ | |

<sup>1</sup> We compare the data of the first replication reported in the manuscript (“original” dataset) to data derived from the “alternative” fROIs (“new” dataset). The first replication was chosen because it had a sufficient number of participants ( $n=13$ ) who listened to the same two stories, namely, (i) and (iii).
<sup>2</sup> For the new dataset, one story was used for defining “alternative” fROIs, and the held out story was used to estimate their ISCs independently of the criteria used to define them. The process was then repeated with the two stories in reversed roles, and the resulting two estimates for each fROI were averaged.
<sup>3</sup> For language (MD) fROIs in the original dataset, one run of the language (MD) localizer was used to define fROIs and the second run was then used to estimate their responses independently of the criteria used to define them. The process was then repeated with the two runs in reversed roles, and the resulting two estimates for each fROI were averaged.

### Correlations of network activity within individuals listening to a story twice

The relatively low ISCs in MD regions could be interpreted in two ways: on the one hand, MD regions might closely track linguistic input but do so in an idiosyncratic fashion across individuals. For example, if different people find different sections of the story difficult to comprehend, they might each recruit their MD network at respectively different times. In this case, MD activity time-courses would be stimulus-locked for each individual but would differ across individuals. Alternatively, activity in the MD regions might not be closely linked to the linguistic input at all. These two interpretations can be distinguished by correlating signal time-courses within a given individual who is listening to the same story twice (Hasson et al., 2009): if MD activity tracks the story in an idiosyncratic manner across individuals, then it should still be similar across two instances of the same story within an individual; however, if MD activity does not track the story, then it should not exhibit reliable time-courses even within an individual.

Therefore, we scanned several participants listening to stories twice and computed within-subject correlations (WSCs). In line with our findings above, WSCs in the LH language network (*r*=0.160) were stronger than in the RH language network (*r*=0.129; *d*=0.33, *z*=3.66, *p*<0.001), the MDfp network (*r*=0.083; *d*=0.83, *z*=8.5, *p≈*0) and the MDco network (*r*=0.097; *d*=1.25, *z*=6.05, *p*<10^−8^). WSCs in the RH language network were stronger than those in the MDfp network (*d*=0.30, *z*=4.48, *p*<10^−4^) and the MDco network (*d*=0.32, *z*=2.66, *p*=0.012), but the two latter networks did not differ (Figure 4a). When we directly contrasted WSCs to ISCs (the latter re-computed as “pairwise-ISCs” to be directly comparable to the former; see Materials and Methods) we found that the patterns of results were indistinguishable across the two measures (for all comparisons between WSCs and pairwise-ISCs, *p*>0.52) (Figure 4b). Therefore, even across story repetitions within a given individual, MD network activity is significantly less reliable than language network activity, indicating that the former, but not the latter, tracks linguistic input closely.

**Figure 4.**
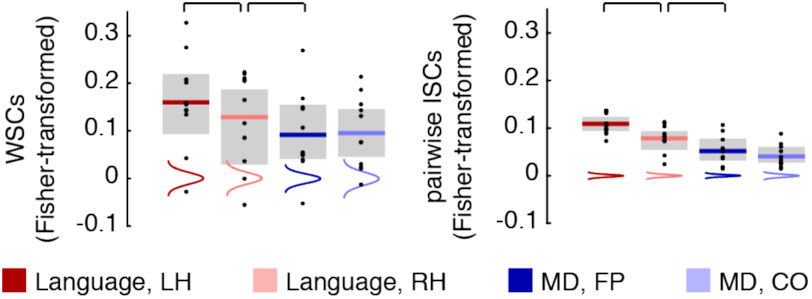
WSCs (left) and pairwise-ISCs (right) during story comprehension in the language and MD networks. Same conventions as in Figure 3.

## Discussion

During story comprehension, a robust and reliable difference in neural activity distinguished between the language network and the MD network. The language network, particularly in the LH, showed relatively little individual variation in activity (high ISCs) due to close tracking of the story (high WSCs). In contrast, MD network activity was more idiosyncratic across individuals (low ISCs), showing weaker tracking of the story (low WSCs). These findings suggest a novel typology of mental processes contributing to language comprehension: it is not only a question of which linguistic features are tracked by different mechanisms, but of whether (and to what extent) these mechanisms track linguistic input. Thus, some processes implemented in the language network are stimulus-related and consistent across individuals; other processes, implemented in the MD network, are less tightly coupled to the input and appear more idiosyncratic.

This distinction importantly constrains cognitive models of language processing: it narrows the space of domain-general processes that can be implemented in the MD network to those processes that do not require continuous access to the input. This conclusion is inconsistent with the assumption of close input tracking, which implicitly underlies existing interpretations of MD network activity in task-based neuroimaging studies of comprehension. It might also be inconsistent with current psycholinguistic models describing how domain-general working-memory resources contribute to incremental, moment-to-moment language processing along with language-specific knowledge (for a review, see: Levy, 2013).

Characterizing the respective contributions of the language and MD networks to comprehension was methodologically possible due to the localization of these networks using functional contrasts, individually for each participant. This method accounts for inter-individual variability in the mapping of function onto cortical anatomy (Jones and Powell, 1970; Gloor, 1997; Amunts et al., 1999; Tomaiuolo et al., 1999; Wise et al., 2001; Chein et al., 2002), conferring high functional resolution (Nieto-Castañón and Fedorenko, 2012) that is unobtainable if regions of interest are instead defined based on anatomical criteria or group analyses of functional data (Juch et al., 2005; Poldrack, 2006; Saxe et al., 2006; Fischl et al., 2008; Tahmasebi et al., 2011; Frost and Goebel, 2012). Consequently, single-participant functional localization provides a principled way of relating our ISC data to known functional divisions in the cortex. This method thus augments the ISC approach, allowing us to provide a novel key characterization of the functional topology of ISCs based on the distinction between the language and MD networks.

Within this topology, the role of MD regions in language comprehension is particularly interesting. Whereas task-based studies have demonstrated that MD regions scale their activity with increasing comprehension difficulty in numerous contexts (Stromswold et al., 1996; Stowe et al., 1998; Caplan et al., 1999; Fiez et al., 1999; Fiebach et al., 2002; Chee et al., 2003; Constable et al., 2004; Rodd et al., 2005; Chen et al., 2006; Nakic et al., 2006; Nieuwland et al., 2007; Novais-Santos et al., 2007; Hauk et al., 2008; Yarkoni et al., 2008; Carreiras et al., 2009; January et al., 2009; Peelle et al., 2009; Ye and Zhou, 2009; Barde et al., 2012; McMillan et al., 2012; McMillan et al., 2013), we demonstrate that they track natural language relatively weakly. One might suggest that the domain-general operations of the MD network are only recruited when linguistic labor is sufficiently high and burdens the language network beyond its capacities; as long as this threshold is not crossed, the working-memory resources that aid in comprehension might be domain-specific and implemented in the language network. However, we find such an interpretation unlikely given that our story stimuli contain frequent occurrences of challenging linguistic phenomena that correlate with behavioral measures of online comprehension difficulty (Shain et al., 2016).

Our finding that the MD network tracks linguistic stimuli relatively weakly also appears to disagree with prior evidence that this network tracks other naturalistic stimuli that are not purely linguistic. Specifically, in audiovisual movies, experiential features like “suspense” modulate MD activity similarly across individuals (Naci et al., 2014), possibly by influencing the frequency of attentional disengagement (Nakano et al., 2013). Does the domain-general MD network play a different role in language comprehension compared to its role in processing other naturalistic stimuli?

Perhaps MD regions are biased towards visual information (or audio-visual integration) in movies compared to the auditory information of stories (Michalka et al., 2015; Braga et al., 2016). Alternatively, MD regions may track both movies and stories, but fluctuations in MD activity during movie viewing could simply be slower, and thus more reliably measured, compared to the fast fluctuations during story comprehension. Therefore, evidence of stimulus tracking by MD regions during story comprehension might only be evident at high frequencies that cannot be measured with the temporally slow BOLD signal of fMRI. Still, we note that the temporal resolution of fMRI was sufficient to capture story tracking in language regions, so the argument above only holds if the MD network tracks stories on a faster time-scale than the language network. Finally, activity in MD regions may reflect internal fluctuations in domain-general attention or “focus” (Norman and Shallice, 1986; Chun et al., 2011) that may co-vary with the emotional manipulations in movies (Williams et al., 2016) but be relatively independent of input processing difficulty during natural language comprehension. This account is also consistent with previous findings of greater MD activity with increased linguistic demands in experimentally designed tasks, insofar as such tasks control the focus of participants more explicitly than naturalistic stories.

To conclude, our study synergistically combines task-based functional localization in individual participants and a naturalistic cognition paradigm for comparing brain activity across participants to characterize the distinct contributions of the language network and MD network to story comprehension. Whereas activity in the language network is similar across individuals and closely tracks stories, activity in the MD network is more idiosyncratic and does not track linguistic input as closely. These findings suggest a novel distinction between different mechanisms that underlie language processing based on individual differences in their processing patterns and their coupling to the linguistic input.

## Acknowledgements

We would like to acknowledge the Athinoula A. Martinos Imaging Center at the McGovern Institute for Brain Research at MIT, and its support team (Steve Shannon, Atsushi Takahashi, and Sheeba Arnold). We thank MIT affiliates Alexander Paunov and Zach Mineroff for their help with data collection, Anastasia Vishnevetsky for her help with constructing the stories and Nancy Kanwisher and Ted Gibson for recording the stories. We thank Uri Hasson (Princeton University) for providing the materials for the second replication. We also thank Nancy Kanwisher, Ted Gibson, John Duncan (Medical Research Council, Cognition and Brain Science Unit), and the audience at the CUNY Sentence Processing conference in San Diego for comments on earlier versions of this work. E.F. was supported by a K99/R00 award HD 057522 from NICHD and by a grant from the Simons Foundation to the Simons Center for the Social Brain at MIT. This work was also partially supported by the Office of the Director of National Intelligence (ODNI), Intelligence Advanced Research Projects Activity (IARPA), via Air Force Research Laboratory (AFRL), under contract FA8650-14-C-7358. The views and conclusions contained herein are those of the authors and should not be interpreted as necessarily representing the official policies or endorsements, either expressed or implied, of ODNI, IARPA, AFRL, or the U.S. Government. The U.S. Government is authorized to reproduce and distribute reprints for Governmental purposes notwithstanding any copyright annotation thereon.

